# Identification and functional analysis of eRNA markers for hepatocellular carcinoma based on high-throughput data

**DOI:** 10.1101/2023.10.28.564513

**Authors:** Zhengxin Chen, Jiaqi Chen, Ruijie Zhang, Yuxi Zhu, Dehua Feng, Huirui Han, Tianyi Li, Xinying Liu, Xuefeng Wang, Zhenzhen Wang, Hongjiu Wang, Limei Wang, Bing Li, Jin Li

**Author notes:** Correspondence: Limei Wang, Bing Li, Jin Li. These authors have contributed equally to this work and share first authorship (Zhengxin Chen, Jiaqi Chen, Ruijie Zhang).

## Abstract

Hepatocellular carcinoma (HCC) is a common type of liver cancer with a high mortality rate. enhancer RNA (eRNA) has been proved to play an important role in cancer progress and development. However, the eRNA studies in HCC are still limited. In this study, we attempted to identify some eRNA biomarkers for HCC diagnosis and analyzed their biological function. First, we identified three eRNA biomarkers (CAP2e, COLEC10e, and MARCOe), which were significant differentially expressed between tumor and normal tissues in 115 HCC patients across three datasets. CAP2e demonstrated upregulation in tumors while COLEC10e and MARCOe were downregulated. These results could be validated in TCGA-LIHC data. There were significant positive correlations between the expression of these eRNAs and their host genes. Then, functional enrichment analysis of protein-coding genes associated with the eRNA biomarkers revealed their involvement in cancer-related pathways. MARCOe was suggested to be a potential target for therapeutic drugs in HCC by a drug related study. The next, survival analysis demonstrated significant prognostic values of these eRNAs in prediction of overall survival. Immune infiltration analysis revealed a positive correlation between MARCOe expression and immune cell infiltration level. Finally, we found similar expression patterns of these eRNA biomarkers in other cancers, such as cholangiocarcinoma, through a pan-cancer comparison. CAP2e and COLEC10e in HCC were validated by other studies. However, the studies about MARCOe in HCC were limited. In conclusion, as best as our knowledge, it is the first time to identify three eRNA biomarkers for HCC diagnosis. These biomarkers are proved to be involved in HCC progress and development, have prognosis prediction values, and are potential to be therapeutic targets.

## INTRODUCTION

Liver cancer, which is the fourth-leading cause of cancer-related death (Villanueva 2019), remains a global health challenge. Its incidence is growing worldwide, with an estimated incidence of more than 1 million cases by 2025 (Llovet et al. 2021). Hepatocellular carcinoma (HCC) is the most common form of liver cancer and accounts for 75% to 85% of cases (Sung et al. 2021). Due to the complex etiology of liver cancer, early diagnosis is rather challenging. By the time most patients receive a definitive diagnosis, the disease has already progressed to an advanced stage (Fu and Wang 2018). Therefore, it is essential to find more effective early diagnostic and prognostic markers to help HCC patients improve their quality of life.

The methods used to diagnose HCC consist of clinical manifestations, imaging techniques (Ayuso et al. 2018), molecular markers (De Stefano et al. 2018), and “Omics” diagnostics (Liu et al. 2019a). In imaging workup, dynamic contrast-enhanced computed tomography and magnetic resonance imaging are deemed the gold standard diagnostic techniques for HCC, while ultrasonography has been extensively utilized as the preliminary screening examination for HCC, demonstrating sensitivities ranging from 51% to 87% and specificities from 80% to 100% (Jiang et al. 2018). Currently, “omics” data such as genomics, proteomics, and metabolomics, have been utilized to detect biomarkers of HCC (Liu et al. 2019a). These diverse biotechnologies have significantly contributed to the detection of HCC biomarkers. Many molecular biomarkers of HCC have been identified, such as alpha-fetoprotein (AFP) (Marrero et al. 2009), Osteopontin (OPN) (Duarte-Salles et al. 2016), and glypican-3 (GPC3) (Scaggiante et al. 2014).

The use of high-throughput sequencing was first proposed in 2005 (Margulies et al. 2005). RNA sequencing (RNA-seq), a key component of high-throughput sequencing, has progressed significantly after more than ten years of development and has become the usual technique for transcriptome research. The range of oncological biomarkers residing in tumor specimens encompass a vast array of DNA, RNA, enzymes, metabolites, transcription factors, and cell surface receptors (Wu and Qu 2015). Groundbreaking evidence has recently illuminated the potential of mRNAs, microRNA (miRNA), long non-coding RNA (lncRNA), and circular RNA (circRNA), as revealed through high-throughput RNA sequencing, to serve as biomarkers for many cancer types (Liu et al. 2019b; Sun and Chen 2020; Zhou et al. 2020; Wang et al. 2022). The enhancer RNAs (eRNAs) are a class of lncRNAs transcribed from enhancer regions of DNA that help regulate the expression of neighboring protein-coding genes. For instance, PSA eRNA is transcribed from an enhancer region of the PSA gene. It can activate AR expression in cancer cells, thereby promoting tumor progression in CRPC (Zhao et al. 2016). The eRNAs are expected to serve as exceptionally useful biomarkers.

Recent studies have systematically documented eRNA expression in humans. The landscape of eRNA expression in cancer tissues was studied using the Cancer Genome Atlas (TCGA) data (Zhang et al. 2019). The analysis of eRNA expression atlas in normal human tissues was performed using the Genotype-Tissue Expression (GTEX) data (Zhang et al. 2021). Several researchers have identified eRNAs as potential targets for therapeutic intervention and valuable biomarkers (Lai et al. 2013; Li et al. 2013; Franco et al. 2015; Rahnamoun et al. 2017; Catarino and Stark 2018). Regarding HCC, Cai et al. utilized the “PreSTIGE” algorithm to identify eRNA associated with immunological genes from TCGA-LIHC (liver hepatocellular carcinoma) data and construct a prognostic model (Cai et al. 2021). Wu et al. identified DCP1A as the most significant survival-associated eRNA from TCGA-LIHC data (Wu et al. 2021). However, there is currently a lack of studies explored differentially expressed eRNAs between normal and cancerous tissues in HCC.

In this study, we identified three eRNA biomarkers for HCC using RNA-seq data and analyzed their functions, prognostic effects, association with immune infiltration and drugs.

## RUSULTS

### Identification of eRNA biomarkers in HCC

By applying McNemar’s Test, seven eRNAs were identified as biomarkers (Figure 1A, B), and their host genes were then identified. Due to the existence of multiple eRNAs corresponding to one host gene, a filtration was performed to select the most important eRNA (tag eRNA) in a gene. In this procedure, the classification accuracy of the eRNA and eRNA combination was used to measure the significance of eRNAs (Supplemental Methods). As a result, three tag eRNAs were identified. ENSR00000194159 exhibited exclusive expression in tumor tissue, while ENSR00000229373 and ENSR00000122322 expressed in normal tissue. The corresponding host genes were CAP2, COLEC10, and MARCO (Table 1).

**Figure 1.**
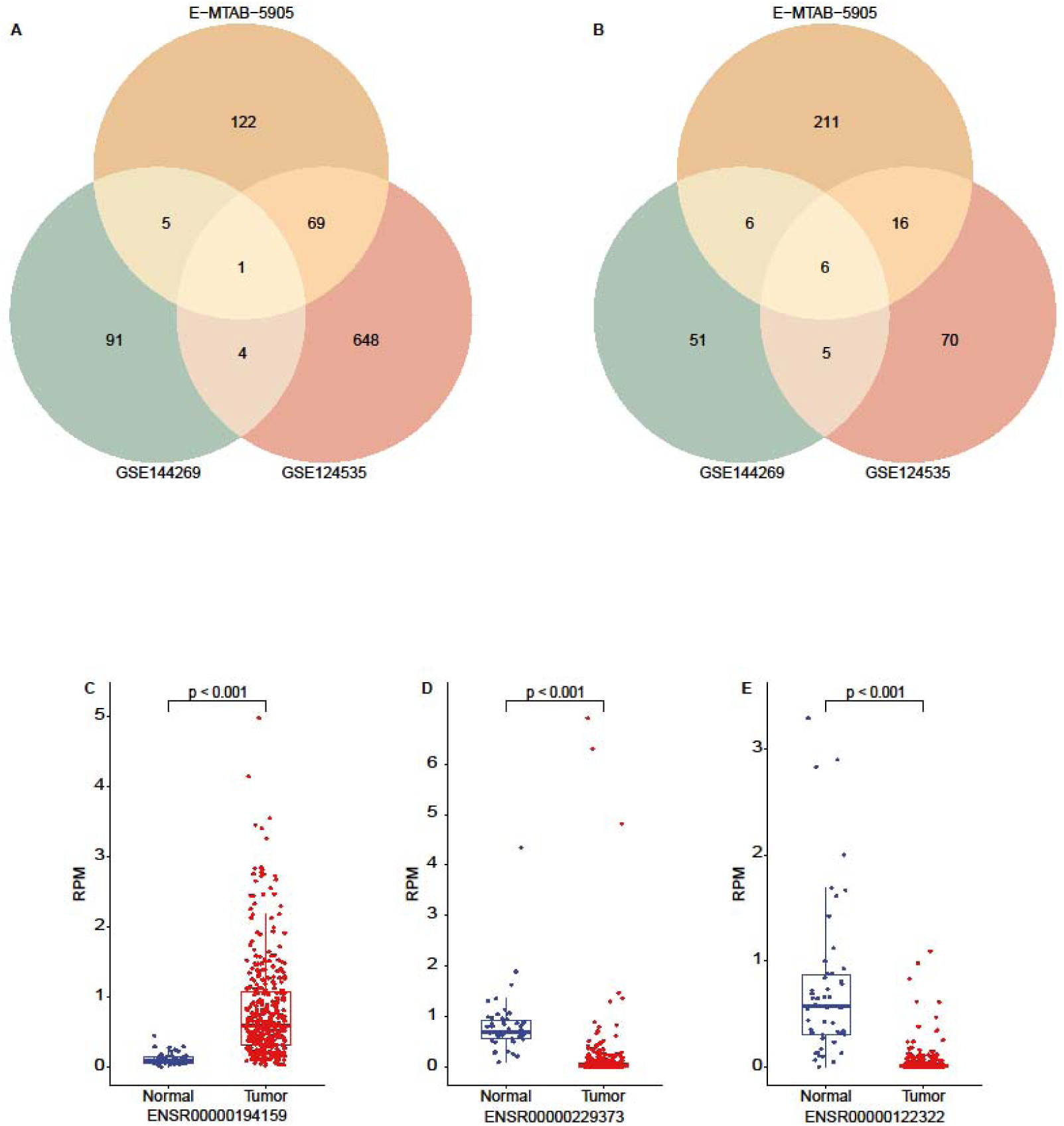
Distribution of eRNA biomarkers. (A) Tumor-Specific eRNA in the multi-datasets. (B) Normal-Specific eRNA in the multi-datasets. (C) The expression levels of ENSR00000194159 in tumor tissues in the LIHC dataset were significantly higher than in normal tissues (P < 0.05). (D) The expression level of ENSR00000229373 in tumor tissues is significantly lower than that in normal tissues (P < 0.05). (E) The expression level of ENSR00000122322 in tumor tissues is significantly lower than that in normal tissues (P < 0.05).

**Table 1.** Tag eRNA biomarkers and the host genes.

| eRNA ID | Chromosome | Start | End | Type | Gene Name | Ensembl ID | Start | End | Distance |
| --- | --- | --- | --- | --- | --- | --- | --- | --- | --- |
| ENSR00000194159 | chr6 | 17522821 | 17528821 | intronic-g | CAP2 | ENSG00000112186.13 | 17393595 | 17557780 | 0 |
| ENSR00000229373 | chr8 | 119080467 | 119086467 | intronic-g | COLEC10 | ENSG00000184374.3 | 118995452 | 119108455 | 0 |
| ENSR00000122322 | chr2 | 118951723 | 118957723 | intronic-g | MARCO | ENSG00000019169.11 | 118942194 | 118994660 | 0 |

These findings were confirmed in the TCGA-LIHC dataset (Fig 1C-E). The expression levels of ENSR00000194159 were significantly upregulated in tumor tissues compared to normal tissues (p < 0.05, Fig 1C). In contrast, the expression levels of ENSR00000229373 and ENSR00000122322 were significantly reduced in tumor tissues (p<0.05, Fig 1D, E).

### Strong correlation between the eRNA biomarker and the host gene

In these three datasets, we utilized Spearman correlation analysis to investigate the relationship between the eRNA biomarkers and their host genes. The results demonstrated a strong positive correlation, with Spearman’s rank correlation coefficient (rho) exceeding 0.8 (p < 0.05, Fig 2 A-I), between the expression levels of ENSR00000194159 (CAP2e) and CAP2, ENSR00000229373 (COLEC10e) and COLEC10, along with ENSR00000122322 (MARCOe) and MARCO. These results emphasize significant consistency between the transcriptional activities of the eRNAs and their host genes, indicating potential interconnected functional roles within the context of HCC.

**Figure 2.**
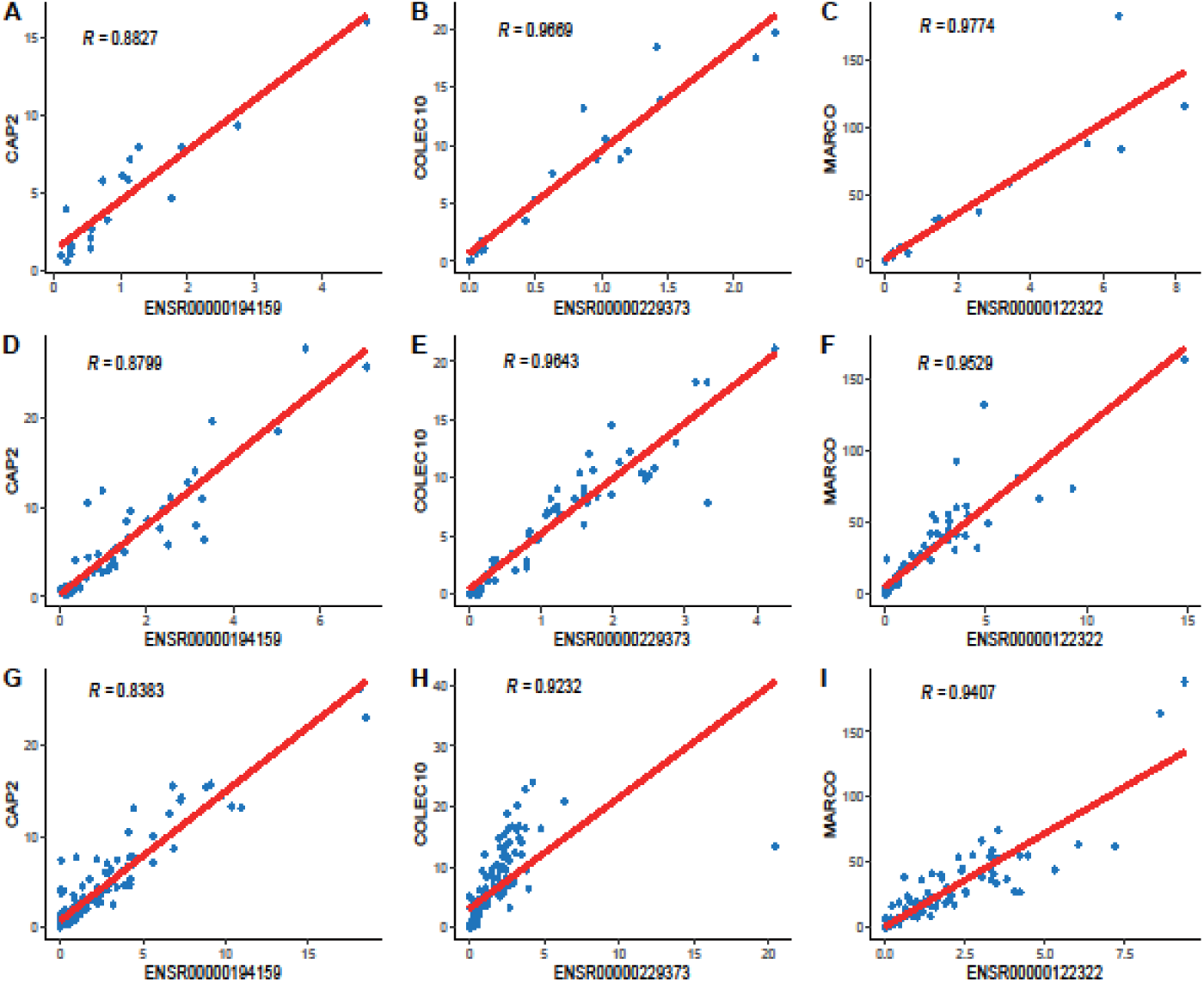
Correlations of expression levels between eRNA biomarkers and host genes across Multiple Datasets. (A-C) Correlation between eRNA biomarkers and host genes in the E-MTAB-5905 dataset; the red line is the fitted curve and “R” is the correlation coefficient. (D-F) Correlation between eRNA biomarkers and host genes in the GSE124535 dataset. (G-I) Correlation between eRNA biomarkers and host genes in the GSE144269 dataset.

### Multiple cancer-related signaling pathways are linked to the eRNA biomarkers

To further explore the potential functions of eRNA biomarkers, Gene Ontology and KEGG Pathway enrichment analysis of protein-coding genes associated with eRNA biomarkers (PCGAeRs) was conducted using DAVID, followed by visualization using R Studio. The results of pathway enrichment analysis indicated significant enrichment in the Cell cycle and Ribosome biogenesis across all three eRNA biomarkers. CAP2e exhibited significant enrichment in several cancer-related pathways, including the Hippo signaling pathway, Glycosylphosphatidylinositol (GPI)-anchor biosynthesis, and Ubiquitin mediated proteolysis (Fig 3A). Similarly, COLEC10e exhibited significant enrichment in several pathways, including Spliceosome, Base excision repair, and N-Glycan biosynthesis (Fig 3B). Finally, MARCOe exhibited significant enrichment in the Basal transcription factors, Base excision repair, and RNA polymerase pathways (Fig 3C). Additional details regarding the functional analysis are discussed in the Supplementary figure S1. Previous studies have shown that hepatitis viral infection, which is the primary promoter of HCC, interacts with ribosomes and plays a major role in the development and advancement of HCC (Xie et al. 2018).

**Figure 3.**
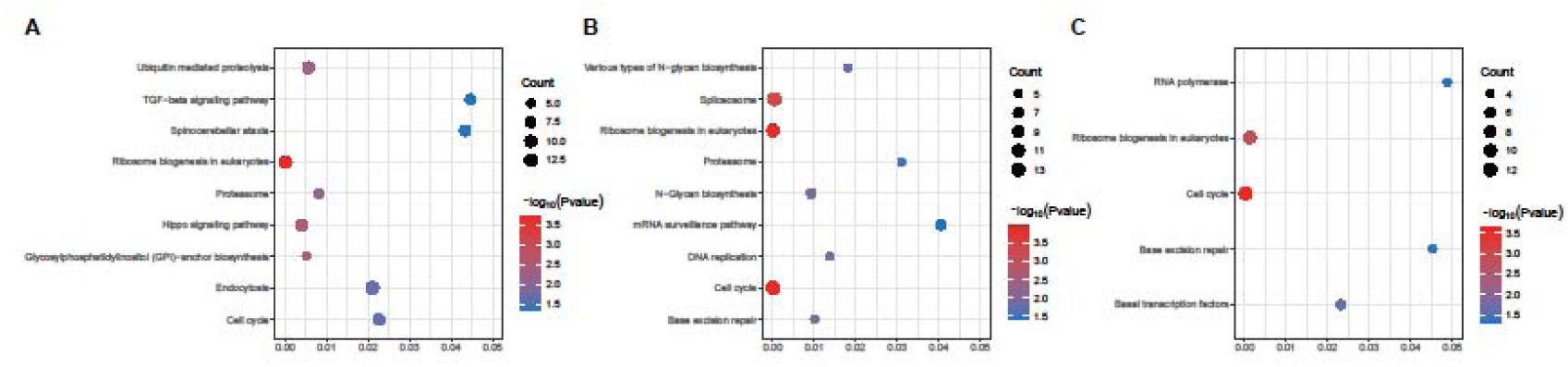
KEGG pathway enrichment analysis of PCGAeR. (A) KEGG pathway enrichment analysis of protein-coding genes associated with CAP2e. (B) KEGG pathway enrichment analysis of protein-coding genes associated with COLEC10e. (C) KEGG pathway enrichment analysis of protein-coding genes associated with MARCOe.

### eRNA is a potential independent prognostic biomarker

To evaluate the potential prognostic prediction of eRNA biomarkers, a survival analysis was performed using the clinical data from TCGA and GSE144269 datasets. Kaplan-Meier analysis revealed that heightened expression of CAP2e exhibited a significant correlation with unfavorable prognosis in both datasets (p<0.05, Fig 4A, D), while decreased expression of COLEC10e was associated with shorter overall survival (p<0.01, Fig 4B, E). Reduced expression of MARCOe was linked to a poor prognosis (p<0.05, Fig 4C) in the GEO dataset, but no significant association was observed in the TCGA dataset (Fig 4F). These findings suggest that eRNA biomarkers hold promise as potential prognostic indicators for HCC.

**Figure 4.**
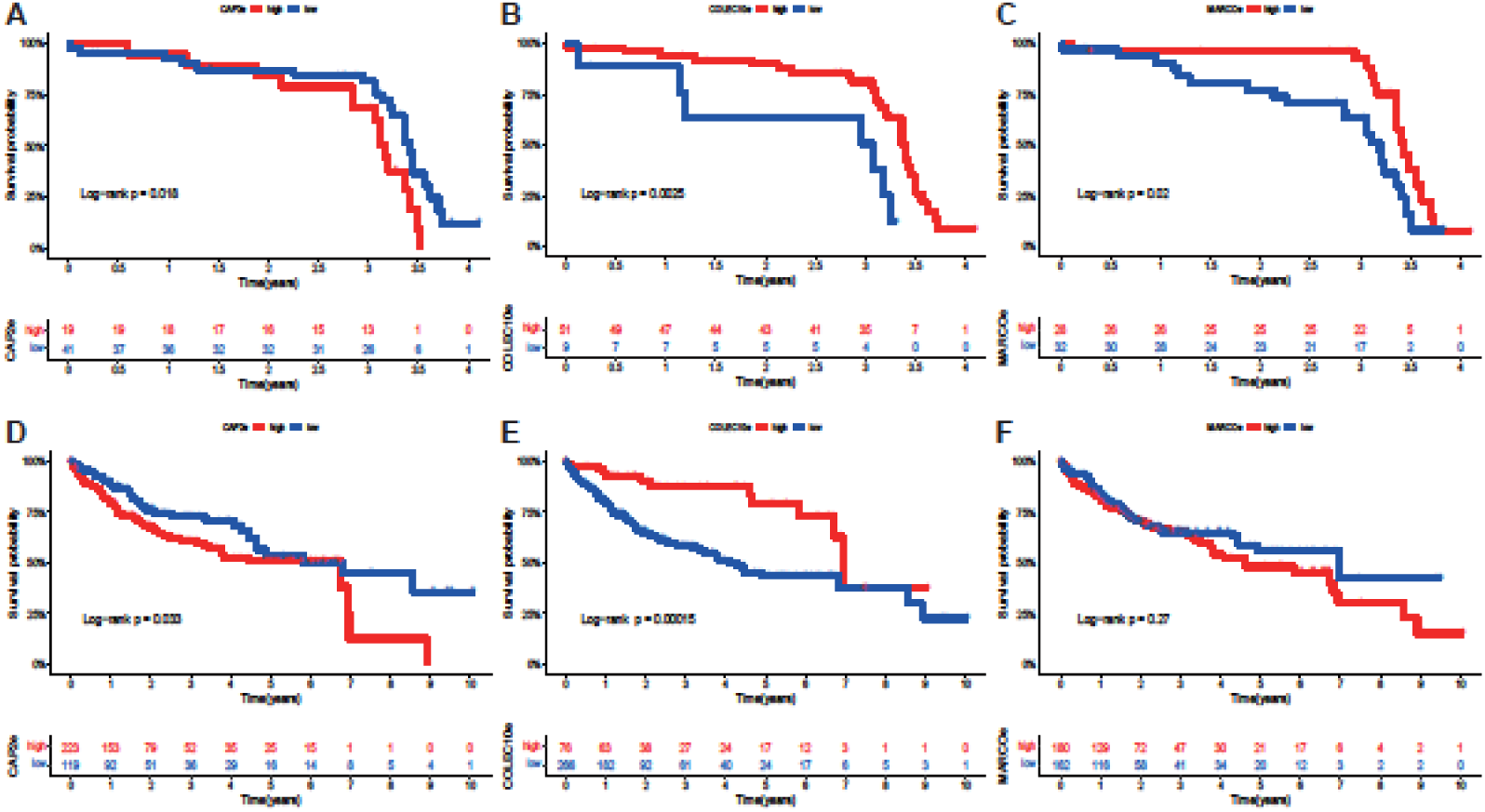
Kaplan-Meier survival analysis was performed on eRNA biomarkers in both the GSE144269 and TCGA-LIHC datasets. (A) The high expression of eRNA CAP2e is associated with poor prognosis in dataset GSE144269. (B) The low expression of eRNA COLEC10e is associated with poor prognosis in dataset GSE144269. (C) The low expression of eRNA MARCOe is associated with poor prognosis in dataset GSE144269. (D) The high expression of eRNA CAP2e is associated with poor prognosis in TCGA. (E) The low expression of eRNA COLEC10e is associated with poor prognosis in TCGA. (F) The expression status of eRNA COLEC10e in relation to prognosis in TCGA.

### The connection between MARCOe and HCC therapeutic medications

The current therapeutic drugs used in the treatment of HCC includes Lenvatinib, Sorafenib, Nivolumab, Cabozantinib, Atezolizumab, Bevacizumab, Imatinib, and others (Ikeda et al. 2018; Tang et al. 2020; Zhao et al. 2020; Liu et al. 2021; Xiao et al. 2021). We obtained their target genes from the HCDT database (Table 2). Then, we conducted pathway enrichment analysis for the top four medications in terms of the amounts of targets (Supplemental Fig S2). The results revealed a significant enrichment of all 4 drugs in the MAPK pathway (FDR < 0.0001). Meanwhile, existing evidence has demonstrated a positive correlation between MARCO expression and the MAPK pathway (Matsushita et al. 2010). Therefore, MARCOe may represent a potential target for therapeutic drugs in HCC.

**Table 2.** Statistics of drug-target number in the HCDT database.

| **Drug name** | **Target**<br>**number** |
| --- | --- |
| Imatinib | 438 |
| Sorafenib | 434 |
| Bevacizumab | 87 |
| Lenvatinib | 33 |
| Cabozantinib | 21 |
| Nivolumab | 16 |
| Atezolizumab | 13 |

**Table 3.** Categorization of McNemar’s Test.

| Normal Tissue | Tumor Tissue |  |
| --- | --- | --- |
| | RPM≥1 | RPM<1 |
| RPM≥1 | A* | B* |
| RPM<1 | C* | D* |

\*A, B, C, and D represent the number of samples in the dataset conforming to the defined criteria.

### Relationship between MARCO expression and immune infiltration

MARCO is an immune-associated gene available in the ImmPort Portal database (Bhattacharya et al. 2014) (https://www.immport.org/home). we utilized Timer 2.0 to examine the association between MARCO expression and immune cell infiltration. MARCO expression was found to have a negative correlation with tumor purity (r = -0.448, p = 1.90e-18, Fig 5). Additionally, MARCO expression was positively associated with the infiltration level of several immune cells, including CD8+ T cells (r = 0.286, p = 6.73e-08), neutrophils (r = 0.245, p = 4.27e-06), macrophages (r = 0.262, p = 8.23e-07), and myeloid dendritic cells (r = 0.317, p = 1.63e-09, Fig 5). Immune infiltration analysis for other eRNA biomarkers is described in the Supplemental fig S3.

**Figure 5.**
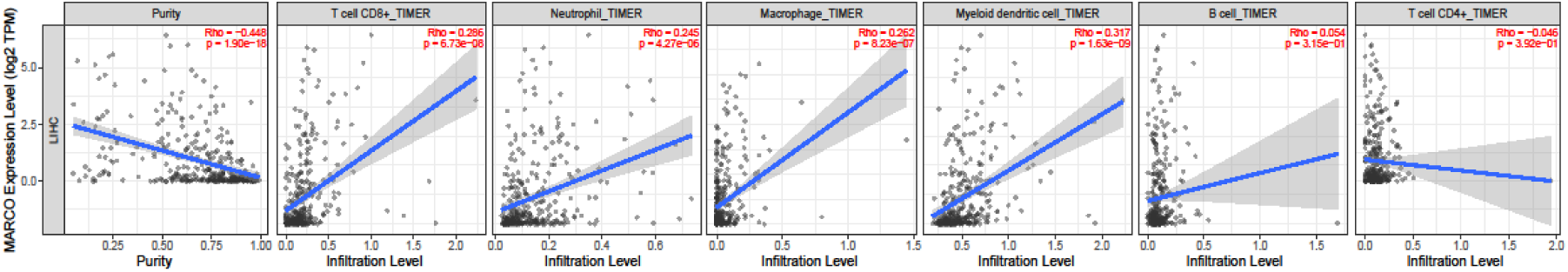
Relationship between the expression level of MARCO and immune cell infiltration

### Pan-cancer analysis of eRNA biomarkers

To investigate the expression patterns of the eRNA biomarker host genes in other cancer types, we performed a pan-cancer expression analysis for CAP2, COLEC10, and MARCO. The findings revealed that these genes exhibited the same pattern in some other cancer types, e.g., CAP2e exhibited a similar expression pattern in CHOL (cholangiocarcinoma) and KIRC (kidney renal clear cell carcinoma) patients as was observed in HCC patients, displaying an upregulated expression in tumor tissues (Fig 6A). Similarly, both COLEC10 and MARCO displayed a comparable expression pattern in CHOL, LUSC (lung squamous cell carcinoma), and LUAD (lung adenocarcinoma) patients as that observed in HCC patients, with downregulated expression in tumor tissues (Fig 6B-C). The shared expression pattern of eRNA biomarkers in CHOL and HCC may potentially be attributed to the differentiation of liver stem cells, which give rise to both hepatocytes and cholangiocytes. Additionally, the phenotypic overlap between LIHC and CHOL has been acknowledged as a continuous spectrum within liver cancer (Kang et al. 2020).

**Figure 6.**
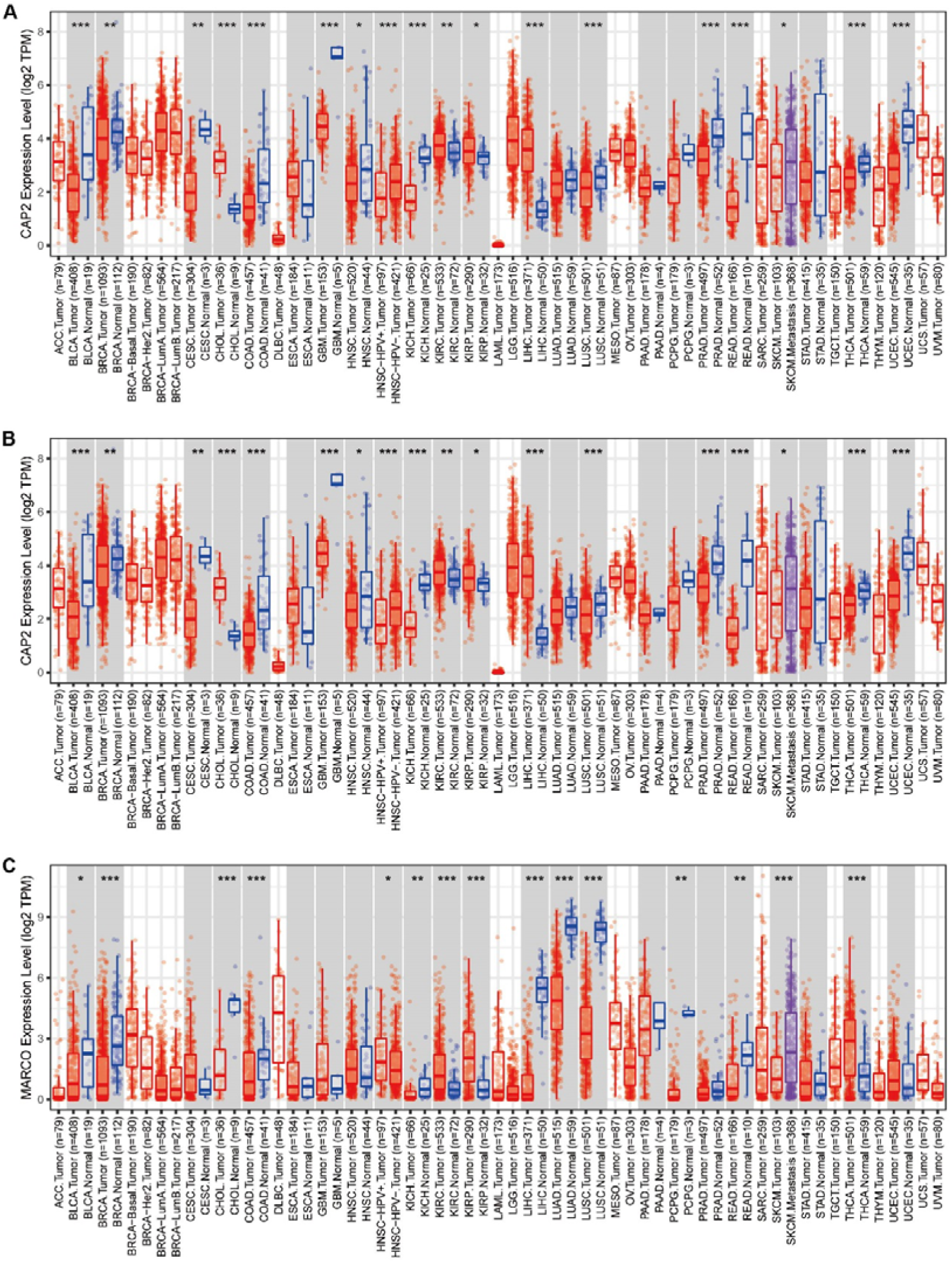
The landscape of host gene expression levels in pan-cancer. (A) The landscape of CAP2 expression levels in pan-cancer. (B) The landscape of COLEC10 expression levels in pan-cancer. (C) The landscape of MARCO expression levels in pan-cancer.

### Literature validation of eRNA biomarkers

Our study unveiled the significant upregulation of CAP2e and its host gene CAP2 in HCC tissues compared to normal tissues. It aligns with Shibata et al.’s research, where they observed elevated levels of CAP2 mRNA and protein in HCC tissues (Shibata et al. 2006). Yoon et al. demonstrated that endoplasmic reticulum stress induces CAP2 overexpression, leading to the activation of Rac1 and ERK, which promote epithelial-mesenchymal transition (EMT) and enhance the migration and invasion of HCC cells (Yoon et al. 2021).

In this study, COLEC10e exhibited significant downregulation in HCC patients and demonstrated significant prognostic value in survival analysis. Ju et al. identified COLEC10 as an immune-related gene (IRG) associated with the tumor microenvironment (TME), and its expression correlated with overall survival (OS) in HCC patients (Ju et al. 2021). Similarly, Zhang et al. discovered that low expression of the COLEC10 encoded protein was significantly associated with vascular invasion and peripheral invasion in HCC patients, establishing COLEC10 as a potential predictive marker for HCC prognosis (Zhang and Wu 2018).

Interestingly, MARCOe exhibited similar biological characteristics to COLEC10e in our study. Previous research has indicated the crucial role of MARCO in the uptake and clearance of tumor cells by macrophages (Xing et al. 2021). A recent study analyzed the correlation between MARCO, immune infiltration, and prognosis in pan-cancers using the data from TCGA and GTEX (Dong et al. 2023). However, studies focusing on MARCO in HCC patients remain limited.

## DISCUSSION

Abnormal eRNA expression has been closely linked to cancer development in various cancers, as numerous studies have shown(Lee et al. 2020; Adhikary et al. 2021). Few researchers have examined eRNAs associated with HCC against normal and cancerous tissues. This study identified three new eRNA biomarkers (CAP2e, COLEC10e, and MARCOe), exhibiting differential expression in HCC tumor tissues relative to normal liver tissues. The results suggest the potential value of these eRNAs as diagnostic and prognostic biomarkers as well as therapeutic targets for HCC.

Our research revealed that tumors tissues showed significantly higher levels of CAP2e than normal tissues, while COLEC10e and MARCOe expression was noticeably lower in tumour tissues compared to normal tissues. The target genes for these biomarkers have been predicted. Subsequently, a correlation analysis was conducted using the expression values of the target genes and eRNA biomarkers; this confirmed the initial prediction as there was a statistically significant positive correlation between them.

Pathway analysis of PCGAeRs showed enrichment in cancer-related pathways for all three eRNAs, emphasizing their broad relevance in HCC pathogenesis. Validation of the prognostic value of CAP2e and COLEC10e supported their clinical usefulness as independent biomarkers. The pan-cancer analysis also exhibited common expression trends for these eRNAs in cholangiocarcinoma, suggesting their involvement in other types of cancer.

Overall, this study presents a systematic process for identifying functionally and clinically relevant eRNA biomarkers from RNA-seq data. The utilization of multiple datasets improves the reliability of the presented results. By clarifying eRNA-mediated gene regulation, new targets, and strategies for the treatment of HCC may be identified.

## METHODS

### Data Collection and Preprocessing

Three RNA-seq datasets were utilized in this study. The high-throughput sequencing data GSE144269 and GSE124535 were downloaded from the GEO database (https://www.ncbi.nlm.nih.gov/geo) (Barrett et al. 2013). The GSE144269 dataset included 140 samples from cancerous tissues and adjacent normal tissues of 70 HCC patients, and the GSE124535 dataset included 70 samples from cancerous tissues and adjacent normal tissues of 35 HCC patients. The third dataset, E-MTAB-5905, was obtained from the EMBL-EBI database (https://www.ebi.ac.uk) (Cook et al. 2020). It included 62 tissues from 11 HCC patients, including cancerous tissues and normal tissues. For each patient, we merged the multiple cancerous tissues belonging to them, as well as the normal tissues. A patient with only one cancerous tissue in this dataset was removed in the following analysis. Finally, we obtained combined eRNA expression level data for 10 patients.

The annotation of enhancers was acquired from Ensembl (https://useast.ensembl.org/) (Zerbino et al. 2015), FANTOM (http://fantom.gsc.riken.jp/index.html) (Andersson et al. 2014), and Roadmap Epigenomics (http://www.roadmapepigenomics.org/) (Bernstein et al. 2010). These three datasets were merged, and only enhancers annotated in a minimum of two datasets were utilized. The eRNA region was defined as the ±LJ3LJkb region surrounding the central locus of the enhancer (Zhang et al. 2019). The human reference genome was collected from the UCSC Genome Browser Home (http://www.genome.ucsc.edu/, hg38) (Karolchik et al. 2011). Alignments of the raw RNA-seq data to the human genome were performed using this reference. Annotation of protein-coding genes were obtained from GENCODE (https://www.gencodegenes.org/, v39) (Wright et al. 2016) and used to obtain the expression levels of protein-coding genes. We downloaded the raw sequence data of LIHC from TCGA data portal (Weinstein et al. 2013) and used it for the validation of eRNA biomarkers. Clinical features for TCGA-LIHC were downloaded from the University of California Santa Cruz (UCSC) Xena Browser built on TCGA data (https://xenabrowser.net) and used for the prognostic analysis of eRNA biomarkers.

During the processing of the raw RNA-seq data, fastp software (Chen et al. 2018) was used to trim the adapter, fastqc software (Andrews 2014) was used to visualize sequence quality, and Hisat2 software (Kim et al. 2019) was used to map the human reference genome (hg38) to clean sequence data. The SAMtools (Li et al. 2009) toolkit was used to convert the file format and merge data from organizations belonging to the same sample in the E-MTAB-5905 dataset. The eRNA annotation file and processed RNA-seq data were implemented as inputs in Bedtools (Quinlan and Hall 2010). The output included the raw read counts for eRNA, which were then converted to Reads Per Million mapped reads (RPM) values in R. Subsequently, for each sample, the eRNA was defined to be present in this sample if its RPM value ≥ 1. StringTie was utilized to derive raw read counts for protein-coding genes from the processed RNA-seq files. The Ballgown package in R was then used to convert the raw read counts into FPKM values (Pertea et al. 2016). A total of 19,982 protein-coding genes were screened in these datasets.

### Research process

After we obtained the RNA-Seq datasets, eRNA annotation, and gene annotation, we got the eRNA and gene expression in 3 datasets. Subsequently, eRNA biomarkers for HCC patients were identified from the eRNA expression matrices, and survival analysis was performed on these eRNA biomarkers using LIHC data from TCGA as validation. Correlation analysis between these eRNA biomarkers and protein-coding genes was conducted to identify protein-coding genes associated with eRNA biomarkers (PCGAeR). Function analysis was then performed for these PCGAeRs. The host genes of these eRNA biomarkers were uncovered and subjected to pan-cancer and immune infiltration analysis. Drug-target information for common chemotherapy drugs in HCC was collected from the HCDT database and integrated with the eRNA-host gene relationships. The flowchart of the entire study is shown in Figure 7.

**Figure 7.**
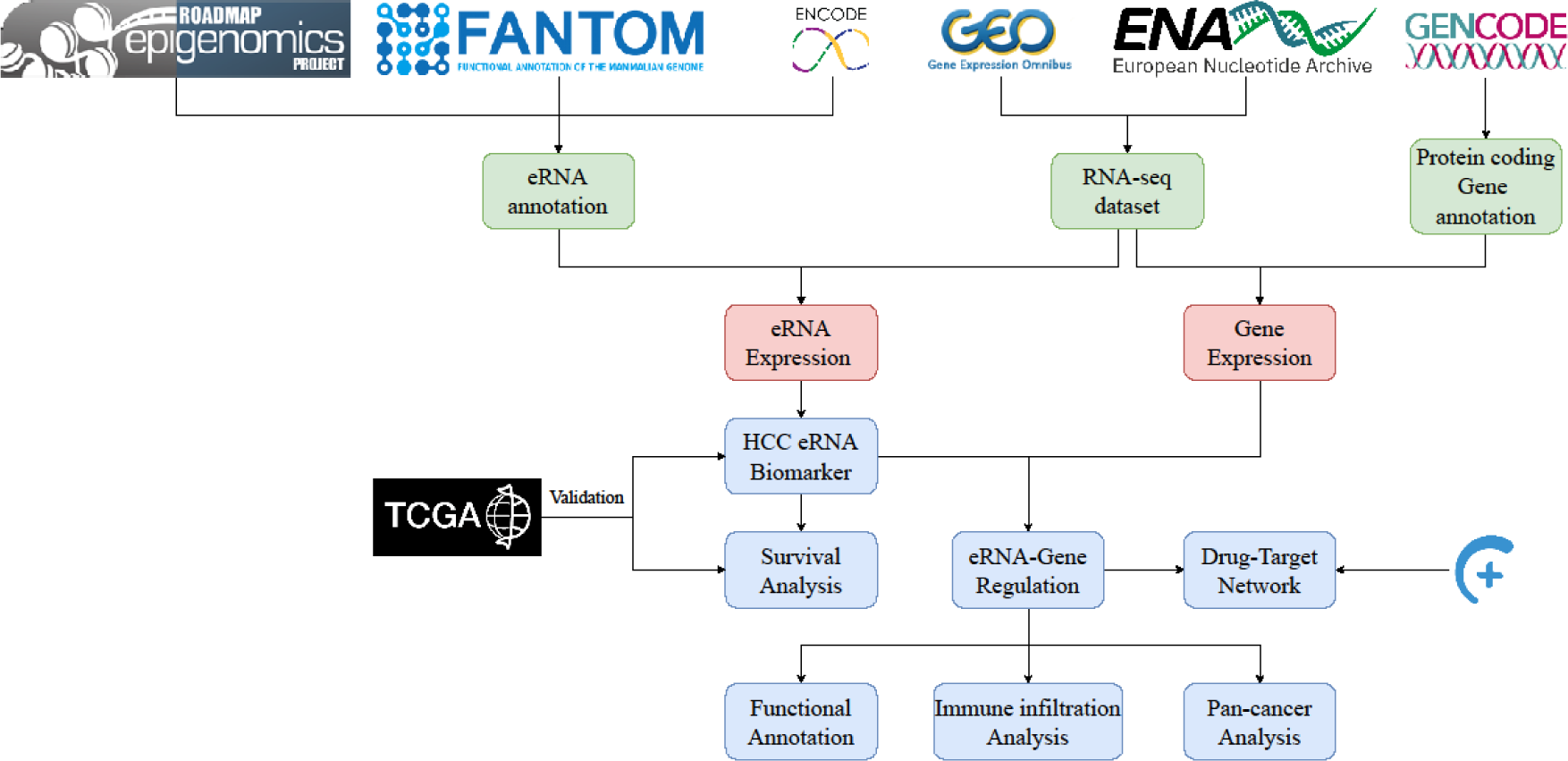
The flow chart of this study

### Screening for eRNA biomarkers and the host genes

For each eRNA in each dataset, we used McNemar’s Test (formula 1) for tumor and normal tissues to identify potential biomarker. P value < 0.05 was considered statistically significant. The categorization and formula were as follows:

Due to the limitations of McNemar’s Test, it cannot cover all the data in the table as A and D were not used. For a useful biomarker, B or C should be large, but the A and D should be small. Therefore, we determined an eRNA to be a biomarker using 3 criteria: It had to be present in all three datasets, have a sample count in group B or C that was larger than or equal to 60% of the overall sample count, and had a significant McNemar’s Test p-value. As a result, seven eRNAs were screened as potential biomarkers (one present in tumor tissue and six in normal tissue). Several recent investigations have shown that eRNAs can function as cis-acting elements (Trapnell et al. 2010; Li et al. 2013; Pefanis et al. 2015; Rahnamoun et al. 2018; Tsai et al. 2018). We predicted the host gene of the eRNA biomarkers based on the closest distance using BEDTools, and ultimately discovered three host genes. The specific patterns of eRNA biomarkers were validated within the TCGA-LIHC dataset.

### Correlation analysis of eRNA biomarkers with protein-coding genes

Spearman correlation analysis was performed between the RPM values of these three eRNA biomarkers (CAP2e, COLEC10e, MARCOe) and the FPKM values of protein-coding genes. The statistical significance was set to be p < 0.05. Protein-coding genes were classified into positively and negatively correlated genes based on the spearman correlation coefficient with the eRNA. The top 1% correlated protein-coding genes were defined as PCGAeRs and the summary statistics were in Table 4.

**Table 4.** Number of protein-coding gene associated with eRNA biomarkers in the datasets.

| eRNA biomarker | E-MTAB-5905 |  | GSE124535 |  | GSE144269 |  | union |  |
| --- | --- | --- | --- | --- | --- | --- | --- | --- |
|  | + | - | + | - | + | - | + | - |
| ENSR00000194159 | 140 | 31 | 145 | 33 | 157 | 31 | 414 | 82 |
| ENSR00000229373 | 49 | 122 | 36 | 141 | 68 | 121 | 99 | 348 |
| ENSR00000122322 | 40 | 131 | 33 | 144 | 68 | 120 | 91 | 354 |

### Gene ontology and KEGG pathway enrichment analysis

To identify the biological functions of PCGAeRs in HCC, we performed functional enrichment analysis for Gene Ontology and KEGG using DAVID (Dennis et al. 2003), and the significance threshold was set at a p-value < 0.05. The results were visualized utilizing the “ggplot2” (v 3.3.6) R package.

### Prognostic analysis of eRNA biomarkers in HCC patients

A survival analysis was performed using GSE144269 and TCGA-LIHC datasets. The samples in GSE144269 and TCGA-LIHC datasets that lacked clinical information were filtered then the samples were split into two groups based on the best cut-off value for eRNA biomarker expression. To determine the cut-off value, the receiver operating characteristic (ROC) was constructed, and the cut-off value was chosen based on the highest sensitivity and smallest distance from the cut-off value to the upper left corner of the ROC curve. Kaplan-Meier survival curves were created using the “survival” (v 3.3.1) R package to discern potential associations between eRNA biomarkers and the prognosis of HCC. The statistically significant threshold was set at a log-rank p-value < 0.05.

### Association of host gene with therapeutic drugs

We collected therapeutic drugs for the treatment of HCC, which were obtained from several papers available on PubMed. Highly Confident Drug-Target Resource (HCDT, http://hainmu-biobigdata.com/hcdt/) was a combined database for drug-target interactions (Chen et al. 2022). We used HCDT to identify the target genes for these therapeutic drugs. KEGG pathways enrichment in DAVID was used to analyze these target genes, then we speculated that the eRNA-Gene-Pathway-Drug relationships.

### Pan-cancer and immune infiltration analysis

The correlation between eRNA biomarkers and the expression levels of immune-infiltrating cells was explored in HCC, and expression levels of eRNA biomarkers were examined in other cancers in the TCGA dataset. TIMER 2.0 was utilized to investigate the expression landscape of host genes for eRNA biomarkers across various cancers in the TCGA dataset and their infiltration level in various types of immune cells (Li et al. 2020).

## DATA ACCESS

The sequencing data for this study is available at the NCBI Gene Expression Omnibus database (GEO; https://www.ncbi.nlm.nih.gov/geo/) under the accession numbers GSE144269 and GSE124535, or at the European Bioinformatics Institute (EBI; https://www.ebi.ac.uk) under the accession number E-MTAB-5905. The code presented in this study are available upon request from the corresponding author.

## ACKNOWLEDGMENTS

This work was supported by the Natural Science Foundation of Hainan Province [No. 621MS041, 821MS045]; National Natural Science Foundation of China [No.32260155].

